# Investigating birds as vectors of the nematode *Litylenchus crenatae* subsp. *mccannii*, the causal agent of beech leaf disease

**DOI:** 10.1101/2025.09.01.673584

**Authors:** Spencer R. Parkinson, Danielle K. H. Martin, Scott H. Stoleson, James B. Kotcon, David J. Burke, Mihail R. Kantor, Christopher M. Lituma

## Abstract

Beech leaf disease (BLD) is an emerging forest disease primarily affecting American beech (*Fagus grandifolia*, Ehrh.) in North America. Tree mortality of sapling-sized trees has been observed within five to seven years of infection. The *Litylenchus crenatae* subsp. *mccannii* (Lcm) nematode is currently recognized as the major causal pathogen of BLD. Scientists first observed BLD in Ohio in 2012 and it has since spread and been detected in 15 states in the USA and one Canadian province. Understanding how this novel pathogen disperses is important for the development of management strategies and predictive models. The spread of BLD to new spatial areas may be due to the transportation of Lcm by animal vectors. We investigated birds as dispersal vectors of Lcm via ectozoochory (external) and endozoochory (internal). The ectozoochory objective was to determine if Lcm can be detected from ectoparasites and or feathers collected from wild birds caught in BLD areas. The endozoochory objectives were to determine if Lcm can be detected in the feces of wild-caught birds in BLD-infected areas, and if viable Lcm can be detected in feces from wild-caught birds kept in cages and fed Lcm nematodes. From 2021 to 2023 we collected 70 ectoparasite and feathers samples (ectozoochory) and 149 fecal samples (endozoochory) from 156 birds in areas in Ohio and Pennsylvania exhibiting BLD. We used molecular and microscopy methods to assess the presence of Lcm. Results confirmed Lcm presence in seven ectozoochory samples, and 14 endozoochory samples from six avian species: tufted titmouse (*Baeolophus bicolor*), black-capped chickadee (*Poecile atricapillus*), white-breasted nuthatch (*Sitta carolinensis*), dark-eyed junco (*Junco hyemalis*), American goldfinch (*Spinus tristis*), and downy woodpecker (*Dryobates pubescens*). These novel findings are among the first to detect plant parasitic nematode DNA in avian biological samples. The methods described in this manuscript could be applied to future BLD research as well as studies on animal vectors and forest pathogens. Understanding which bird species are vectors of Lcm will facilitate the creation of predictive models and develop preventative management strategies to protect uninfected American beech stands.

## Introduction

American beech (*Fagus grandifolia*) is a hard-masting, canopy-dominant, hardwood tree of eastern North America. American beech is long-lived, surviving up to 400 years, and typically produces mast beginning around 40 years or 13–20 cm diameter at breast height [1]. American beech serves as the dominant hard-masting species in maple-beech northern hardwood forests and produces a significant mast crop every other year subsequently followed by a poor mast crop. American beech nuts are a food source for >30 species of birds and mammals and alternating masting years directly affect wildlife populations [2,3]. In Maine, during poor mast years, black bear (*Ursus americana*) reproduction decreased by 22%, and American marten (*Martes americana*) trapping was significantly reduced [4]. The introduction of beech bark disease (*Cryptococcus fagisuga and Neonectria spp.*) increased pressure on wildlife populations that utilize American beech nuts and for nesting and foraging. Beech bark disease primarily affects mature canopy-dominant trees, stimulating root sprouting in diseased and dying American beech trees [5]. This results in the replacement of canopy-dominant overstory trees with understory American beech thickets [6]. The transition of canopy trees to American beech thickets altered the biodiversity of northern hardwood forests by suppressing native hardwood seedling growth, specifically sugar maple (*Acer saccharum*) and yellow birch (*Betula alleghaniensis*), and outcompeting other understory plants due to heavy shading of forest floor [7]. The aftermath of beech bark disease left forests structured for a new pathogen outbreak [5].

Beech leaf disease (hereafter: BLD) is a novel forest disease that primarily affects American beech in North America [8]. Scientists first recognized BLD symptoms in Lake County, Ohio in 2012 and BLD has since been detected throughout the Northeastern United States and Southeastern Canada, from Michigan to the Atlantic and from Ontario to Virginia [9,10]. *Litylenchus crenatae* subsp. *mccannii*, (hereafter: Lcm, family Anguinidae) is a recently discovered plant pathogenic nematode highly associated with BLD [8]. In a 2019 study, Lcm was present in 45–90% of leaves with visible BLD symptoms and 90% of buds from symptomatic trees [11]. Scientists described the nematode *Litylenchus crenatae* in Japan in 2019, and Lcm is considered a subspecies in North America, potentially indicating a country of origin [12]. BLD symptoms have also been observed on introduced species of beech in North America: European beech (*F. sylvatica*), Chinese beech (*F. engleriana*), and oriental beech (*F. orientalis*) but not on Japanese beech (*F. crenata*) [11]. Symptoms of BLD include darkening green stripes or bands between lateral veins on leaves in the early stages progressing to thickened, crinkled, leathery, and puckering of leaves with heavy chlorotic striping. Interveinal leaf banding is directly influenced by the Lcm nematode [13]. Recent research by Vieira et al. (2023) has shown that as infected leaves mature, Lcm migrates from the surface to the inner tissues of the leaf, leading to the establishment of populations within spongy mesophyll layers. They also showed that symptomatic leaves infected with Lcm have thicker mesophyll layers, accelerated rates of cell division, and increased chloroplast content compared to controls. Scientists have found several plant pathogens induce the formation of enlarged cells, stimulating a high metabolic activity to provide the parasite with nutrients while depriving the host [13]. However, understanding the rapid spread and distribution of a nematode with limited dispersal ability to remote and disjunct areas is of critical importance to the survival of American beech.

Birds can act as vectors for the long-distance dispersal of human, animal, and plant pathogens [14]. The two methods of dispersal are ectozoochory (external) and endozoochory (internal) [15-17]. Ectozoochoric dispersal occurs when birds collect pathogens on their feet, feathers, or beaks and shed them into uninfected areas. Birds can spread Lyme disease by transporting ticks that carry *Borrelia burgdorferi* bacteria [18-20]. Canada geese (*Branta canadensis*) and graylag geese (*Anser anser*) can spread Chytridiomycosis, an amphibian disease, by transporting chytrid fungus (*Batrachochytrium dendrobatidis*) on their feet [21-23]. Endozoochoric dispersal occurs internally by consumption and successful excretion of pathogens. European starlings (*Sturnus vulgaris*), northern bobwhite (*Colinus virginianus*), domestic rock pigeons (*Columba livia*), wood thrush (*Hylocichla mustelina*), brown-headed cowbird (*Molothrus ater*), and white-throated sparrow (*Zonotrichia albicollis*) can consume potato cyst (*Globodera rostochiensis*), and soybean cyst nematodes (*Heterodera glycines*) and excrete viable egg-bearing cysts between 30 minutes and 48 hours of consumption [24,25].

Birds may be spreading BLD through ectozoochory or endozoochory of Lcm. Lcm is a foliar nematode that could attach to the feathers or legs of birds as they move through dense American beech stands, or as they feed or nest on branches (ectozoochory). Additionally, common avian ectoparasites, such as mites, may be covered in Lcm and subsequently attach to birds. Lcm nematodes and eggs are concentrated in American beech buds in the winter, and birds may consume these buds when other food sources are scarce [26]. Lcm is most concentrated in the buds of infected trees in October [8,27]. Birds may ingest Lcm nematodes through buds and excrete viable nematodes into uninfected American beech areas (endozoochory). Lcm is tolerant of acidic environments, such as a bird’s stomach. In a 2021 experiment, Lcm nematodes subjected to varying pH levels demonstrated resilience; 75% survived a pH of 2, and 10% survived a pH of 1.7 after 24 hours [28]. This pH level aligns with the acidity of a vulture’s stomach (Family Cathartidae), suggesting that Lcm could endure the higher pH stomachs of forest birds [29]. Lcm DNA can be extracted and detected with PCR in chicken feces inoculated with Lcm in a lab setting [30]. We hypothesized that common forest birds can serve as dispersal vectors of Lcm through ectozoochory or endozoochory. Our objectives for ectozoochory were to determine 1) if Lcm can be detected from ectoparasites collected from wild birds caught in BLD areas, and 2) if Lcm can be detected in feather samples collected from wild birds caught in BLD areas. We hypothesized that Lcm will be detected in both ectoparasite and feather samples. Our objectives for endozoochory were to determine 1) if Lcm can be detected in the feces of wild-caught birds in BLD-infected areas, and 2) if Lcm can be detected in feces from wild-caught birds kept in cages and fed Lcm nematodes. We hypothesized that Lcm will be detected in the feces of wild-caught birds, and caged birds fed Lcm will excrete live, viable nematodes.

## Materials and Methods

### Study Sites

#### Holden Arboretum

We collected samples at the Holden Arboretum (Holden) in Northeastern Ohio located in Lake and Geauga counties on the High Allegheny Plateau. Our sample sites were Arbor Vitae (41.61061, -81.29452) and Wisner (41.60685, -81.28819, Fig 1A). Arbor Vitae was adjacent to BLD-infested stands near a collection of arborvitaes. Wisner was directly in BLD-infested trees along Wisner Road running along the east branch of the Chagrin River. We sampled in 2022 on January 4th – 6th, April 28th – 29th, August 2nd – 4th; and in 2023 on February 2nd and April 6th. BLD was first observed at Holden in 2014 (the initial detection) and has since spread throughout the arboretum and natural areas infecting American beech, European beech (*F. sylvatica*), Chinese beech (*F. engleriana*), and Oriental beech (*F. orientalis*) [11].

**Fig 1.**
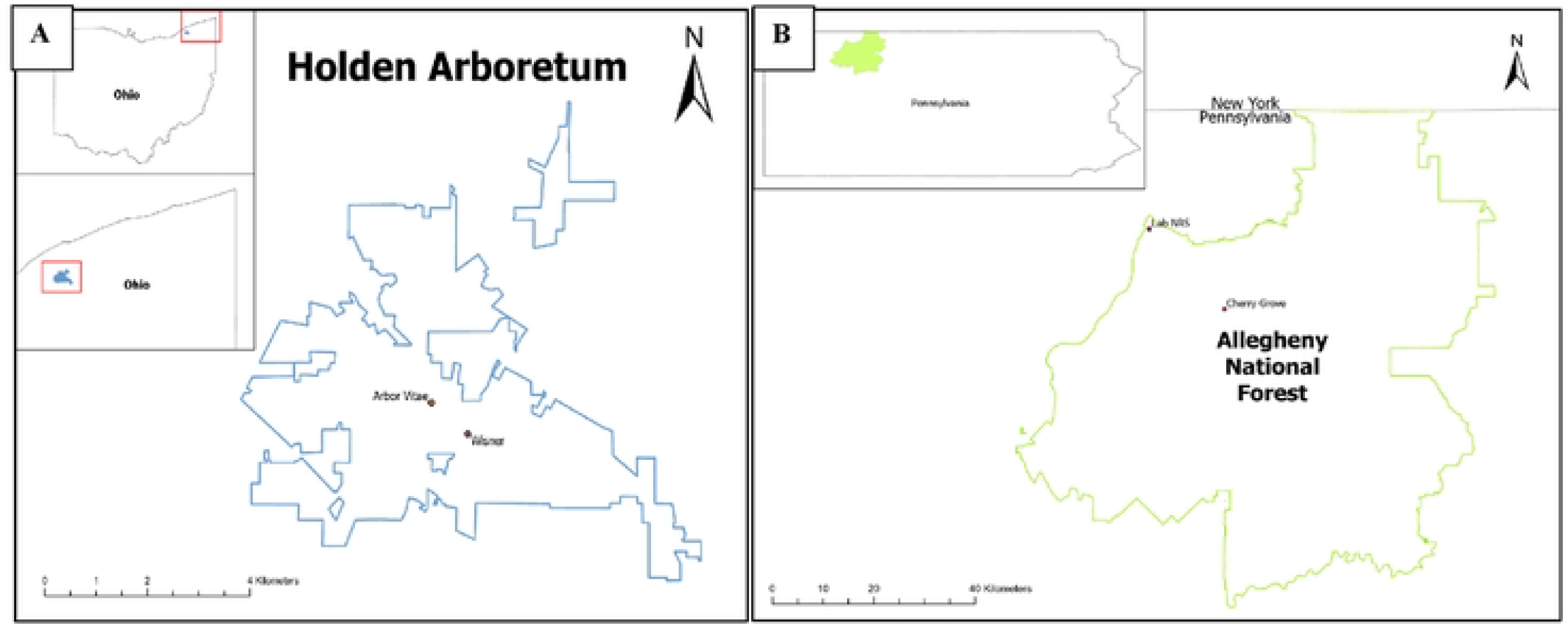
Field sites at the Holden Arboretum and Allegheny National Forest. Map of sites where we caught wild birds, collected American beech buds, and exposed wild birds in cages to buds from 2021-2023 to determine if birds can act as vectors of Lcm. We collected data at (A) Holden Arboretum and (B) Allegheny National Forest, USA.

#### Allegheny National Forest

We collected samples at the Allegheny National Forest (ANF) in Northwestern Pennsylvania in Elk, Forest, McKean, and Warren counties on the Western Allegheny Plateau. Our sample sites were Cherry Grove (41.730124, -79.124289) and Lab NRS (41.83827, - 79.26093, Fig 1B). Cherry Grove was directly in BLD-infested trees at the intersection of Cherry Grove Rd and Forest Road 534. Lab NRS was adjacent to BLD-infested stands on the lawn of the Northern Research Station laboratory. We sampled in 2021 on October 7th; and in 2022 on March 16th – 17th, August 9th, November 2nd – 3rd, and December 1st. Beech bark disease first appeared at the ANF in the early 1980s and by 2010 the killing front covered the entire ANF. The killing front resulted in 60–70% mortality and typically affected larger trees (> 20 cm diameter at breast height [31]). Dense American beech thickets sprouted from the root systems in the wake of beech bark disease. American beech thickets prevent the regeneration of other hardwoods and conifers and are highly susceptible to BLD. BLD is currently found in almost every part of the ANF.

### Avian Sampling Methods

#### Bird Feeder Establishment

We chose sites where American beech was the dominant species, consisting of >50% basal area, and >75% of American beech had visible BLD symptoms. In fall 2021, we established 2-3 Squirrel Buster® bird feeders per study site in areas with BLD symptoms. We set up feeders at least 250 m apart, 2 m above the ground, and filled them with a mixed general bird seed every week. Bird feeders were hung from a metal shepherd’s hook and a motion-activated camera trap (Bushnell 30MP CORE Trail Camera) was attached to a nearby tree that took a single photo at 30-second intervals to confirm bird activity.

#### Bud Collection

We collected buds from BLD symptomatic American beech branches 1-2 days before avian sampling. We confirmed the presence of live Lcm nematodes from buds by soaking 10% of buds in water and observing under a microscope as described below. The remaining buds were stored at 4°C until we were ready to use them in the field. One to two days before avian sampling, we emptied the bird seed from the feeders and filled the feeders with remaining buds.

#### Avian Sampling

We used passive mist nets around bird feeders to catch and collect samples from birds [32]. We used two standard mist nets (6 or 12 m long, 2.6 m high, 36 mm mesh) to catch birds [33]. Mist nets were set up in an L shape at least 4 m from the feeders. Nets were set up 30 minutes before sunrise and kept open for at least 4 hours (weather permitting). We banded all birds with a unique USGS aluminum band. Morphometric data on sex (Male/Female/Unknown), mass (g), and wing chord and tail length (mm) was recorded [34]. Species names were recorded using the American Ornithological Society (AOS) 4-letter alpha code [35]. We followed all institutional, state, and federal guidelines for the care and use of animals and conducted work under Institutional Animal Care and Use Committee protocol # 2108045949.1, US Geological Survey banding permit # 23227, and Ohio Division of Wildlife Scientific Collection License # SC230010.

#### Feather and ectoparasite sampling

We collected feather samples from a subsample of birds using sterile forceps and plucking from the base of the feather shaft [36]. We collected three to five breast feathers, one to two from the crown, and one to two from the vent. Feathers were placed in a sterile 1.5-mL microcentrifuge tube and stored at 4°C until we conducted lab work. For ectoparasite sampling, we used dust-ruffling methods [37] but used diatomaceous earth instead of pyrethrin as a safer alternative for humans and birds. A Crayola size 6 paint brush was used to apply diatomaceous earth to the bird’s breast, vent, and wings taking care to avoid the head and eyes. We brushed diatomaceous earth through feather tracks and onto brightly colored paper. The dust was inspected with a 4x hand lens to look for ectoparasites and collected in a sterile 1.5 mL centrifuge tube for molecular testing. We used molecular and microscopy methods to test for the presence of Lcm DNA in feathers and ectoparasite samples.

#### Bagged bird fecal sampling

We collected fecal samples during all sampling periods from October 7th, 2021 to August 4th, 2023. Birds were placed in a brown paper lunch bag and clipped shut. Birds were kept in bags for no more than 15 minutes regardless of defecation and released [38]. Fecal matter was scraped out of bags using sterile forceps and a scalpel and placed into a sterile 1.5 mL capped micro-centrifuge tube. Samples were kept at 4°C until we conducted lab work.

### Caged Birds

#### Building cages

We used 19-gauge galvanized steel wire mesh to create 20.3 × 20.3 × 25.4 cm bird cages (Fig 2). Two small plastic bird feeder dishes were placed near perches, one for water and one for American beech buds from BLD trees. The cages were built larger than 15 × 15 × 15 cm, the recommended minimum cage size for all the species of birds we sampled [39]. The cages were placed in a 27.9 × 34.3 cm plastic dish pan to collect fecal matter. Between every bird, we cleaned the cages and plastic bins with a 10% bleach solution and a firm dish brush and thoroughly sprayed them with water to avoid cross-contamination.

**Fig 2.**
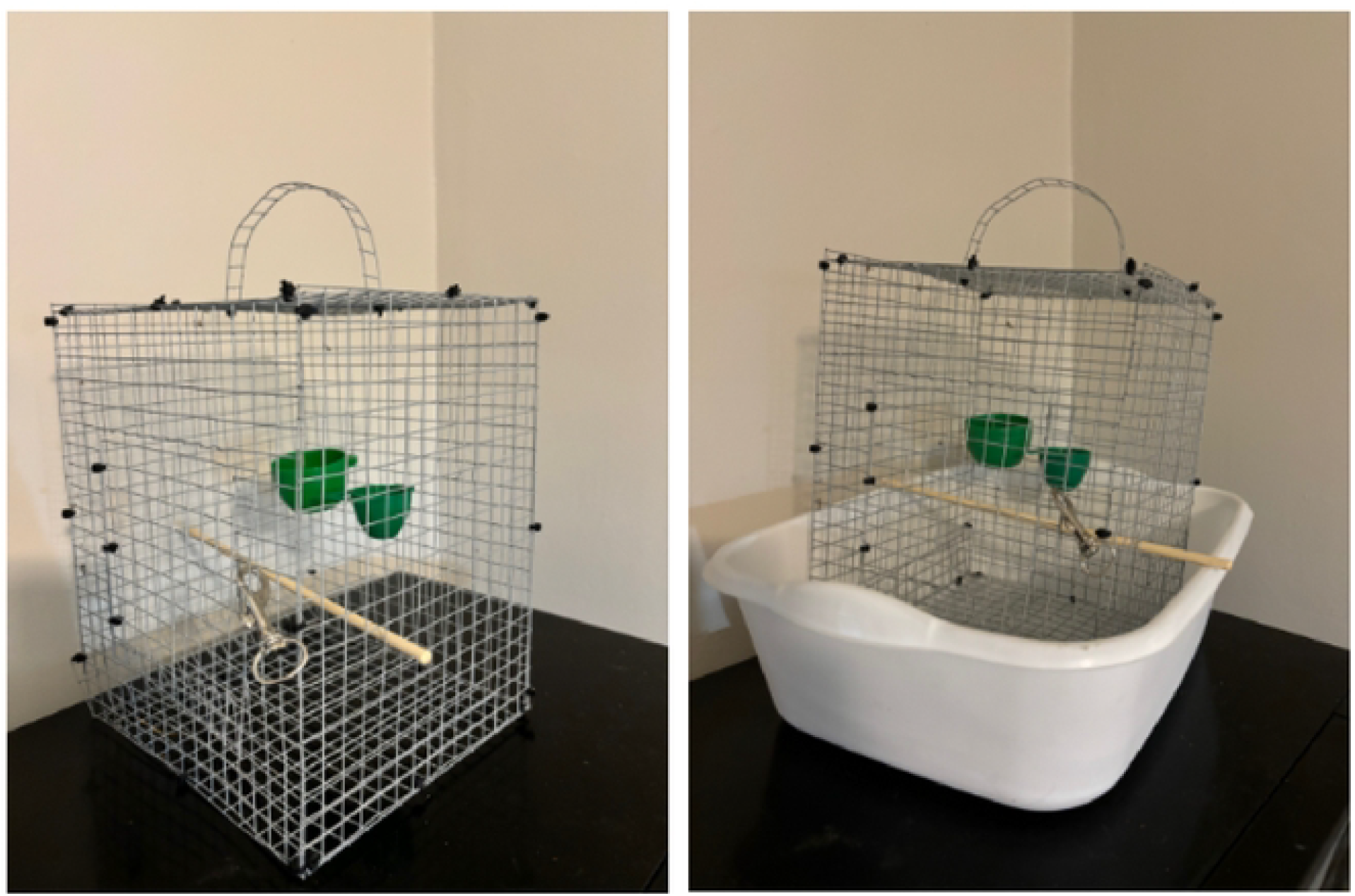
Pictures of Cages. 20.3 × 20.3 × 25.4 cm bird cages made from 0.9 × 15.2 m, 1.27 cm2 19-gauge galvanized steel with plastic bins to collect fecal matter.

#### Feeding nematodes to caged birds

We conducted a caged experiment to ensure birds were consuming American beech buds with BLD. We caught birds at feeders and after processing, if we deemed the birds healthy according to historically reported morphometric measurements and visual examination, we placed birds into an individual cage [34,40]. We prioritized birds based on species that had previously been observed at bird feeders filled with American beech buds: house finch (*Haemorhous mexicanus*), purple finch (*H. purpureus*), tufted titmouse (*Baeolophus bicolor*), chickadee spp. (*Poecile* spp.), and downy woodpecker (*Dryobates pubescens*) (Martin, per comm.). While in cages, birds had access to water and 2 g of BLD-infested American beech buds. Birds were fed two to three buds every 45 minutes. Since we could not 100% confirm that the buds we were feeding the birds contained nematodes we switched to pipetting Lcm nematodes in the birds’ mouths. We extracted live nematodes from infected buds and created a suspension. We pipetted 500-µL of Lcm nematode suspension into each bird’s mouth every 45 minutes. All fecal matter was collected from the plastic floor and placed in a separate sterile 1.5-mL microcentrifuge tube.

Birds were released if they showed signs of stress such as lethargy, crouching, blinking, bleeding, or fluffing [41]. Birds were kept for no more than three hours, which corresponds to the evacuation time for potato cyst nematodes fed to birds [25]. Birds were released at the capture site at least one hour before sunset.

### Laboratory Methods

#### Live nematode collection

To extract live nematodes, we collected leaves and buds from American beech trees with BLD symptoms at the ANF and Holden. We used Baermann funnel and water-soaking methods to collect live Lcm nematodes from plant material [42,43]. For the Baermann funnel extraction, we used three to five leaves or buds. Leaves were cut into 2-cm^2^ pieces, and buds were macerated and placed into a mesh screen. We soaked the mesh screen with plant material in tap water for 24 hours in a 250-mL glass funnel with a 10-cm long section of 1-cm diameter rubber tubing. A binder clip was attached to the bottom of the tubing to keep water from escaping. After 24 hours we released water into a 10-mL test tube and allowed it to settle for at least another hour. The top 9 mL of water was aspirated leaving 1 mL of live nematode suspension at the bottom. For water-soaking extraction, we cut three to five leaves into 2-cm^2^ pieces, or macerated three to five buds and placed them in a 100-mm petri dish. We soaked plant material for 24 hours in distilled water. Plant material was removed from the petri dishes and water was pipetted into 10-mL test tubes and allowed to settle for at least another hour. The top 9 mL of water was aspirated leaving 1 mL of live nematode suspension. We used microscopy to confirm the presence of live nematodes in suspension.

#### Microscopy Methods

For microscopy, we used a 10× to 50× binocular stereoscope to detect live nematodes. To confirm live motile Lcm nematodes from buds and leaves we observed 1 mL of suspension on a 60-mm petri. To feed nematodes to birds, we parsed out nematodes into separate 500-uL suspensions with at least 100 live nematodes present. We looked for Lcm nematodes in the fecal samples by diluting fecal matter with equal parts distilled water and mixing vigorously to create a slurry. The slurry was pipetted onto a 60-mm petri dish and observed with a microscope. Forceps were used to break apart any solids in the feces, if no nematodes were present, we discarded the samples. To look for Lcm nematodes in feather samples we placed feathers in a petri dish and used forceps to tease apart feather barbs. For ectoparasite samples, we used a 10x hand lens to look through dust for ectoparasites.

#### Molecular Methods

For molecular testing, we collected samples in or transferred samples to, a bead-beating PowerBead Tube [Qiagen] with 1 mL of InhibitEX Buffer and vortexed continuously for 1 min to properly break apart samples. DNA was extracted from samples using a QIAamp® Fast DNA Stool Mini Kit [44]. We used a species-specific primer for *Litylenchus crenatae* with forward primer 33F: GGAAAAGGAGCTGACTGGGC; and reverse primer 234R: ACGGACGCAGGCTGTAAGCC [45]. PCR conditions followed protocol from Burke et al. [45], we used a positive and negative control during all PCR amplifications. PCR products were visualized using a standard 1% agarose gel. Positive Lcm amplifications were cleaned up using Wizard® SV Gel and PCR Clean-up System [Promega] and sent to Eurofins Genomics for sequencing. Sequences were aligned using Molecular Evolutionary Genetics Analysis (MEGA) software [46]. We used the basic local alignment search tool (BLAST) with the National Center for Biotechnology Information (NCBI) database to confirm sequenced DNA matched Lcm by at least 99% identity.

## Results

We detected Lcm DNA in 21 samples from 20 individuals. Seven samples from ectozoochory tested positive and fourteen samples tested positive from endozoochory (Table 1). We collected a total of 219 samples from 156 individuals. At the ANF we caught 45 birds (1.2/effort hour). At Holden, we caught 111 birds (2.8/effort hour). We caught nineteen species with black-capped chickadee being the most prevalent, followed by tufted titmouse, dark-eyed junco, American goldfinch, and white-breasted nuthatch (Fig 3).

**Table 1.** DNA detection results by experiment type.

| Study | N | Positives | % Positive |
| --- | --- | --- | --- |
| ENDOZOOCHORY |  |  |  |
| Feces (Bagged Birds) | 90 | 4 | 4.4 % |
| Feces (Caged Birds) | 59 | 10 | 16.9 % |
| ECTOZOOCHORY |  |  |  |
| Ectoparasites | 55 | 6 | 10.9 % |
| Feather Samples | 15 | 1 | 6.7 % |
| Total | 219 | 21 | 9.6 % |

**Fig 3.**
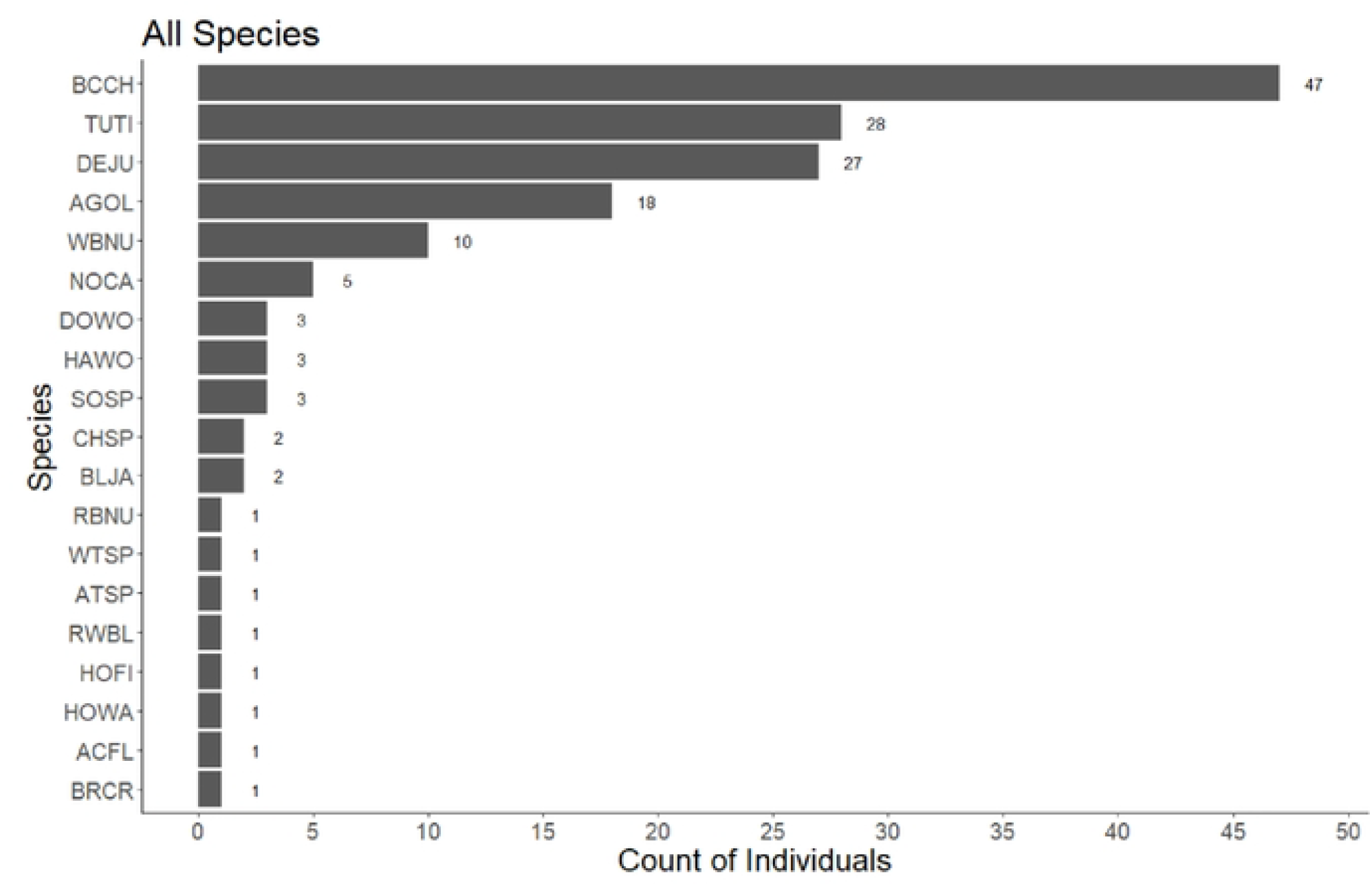
Count of species caught and sampled. Total number of birds caught across all sampling dates and at the ANF and Holden. We used 4-letter alpha codes for bird species. Top five bird species were black-capped chickadee, tufted titmouse, dark-eyed junco, American goldfinch, and white-breasted nuthatch. For all codes refer to: (https://www.birdpop.org/docs/misc/Alpha_codes_eng.pdf).

Samples size, positives, and percent positives for Lcm DNA detected in avian samples for endozoochory and ectozoochory based on different methodologies. Bagged birds were caught and kept in paper bags for < 15 minutes. Caged birds were caught and kept in cages for < 4 hours and fed Lcm nematodes. Ectoparasites were the result of brushing diatomaceous earth through bird feather tracks. Feather samples were feathers plucked from the head, breast, and vent before any other collection was done

### Feather samples

We collected 15 feather samples from birds. One sample, a feather sample from a tufted titmouse from the ANF, was positive for Lcm DNA (6.67%, Fig 4a). We observed no motile nematodes with microscopy.

**Fig 4.**
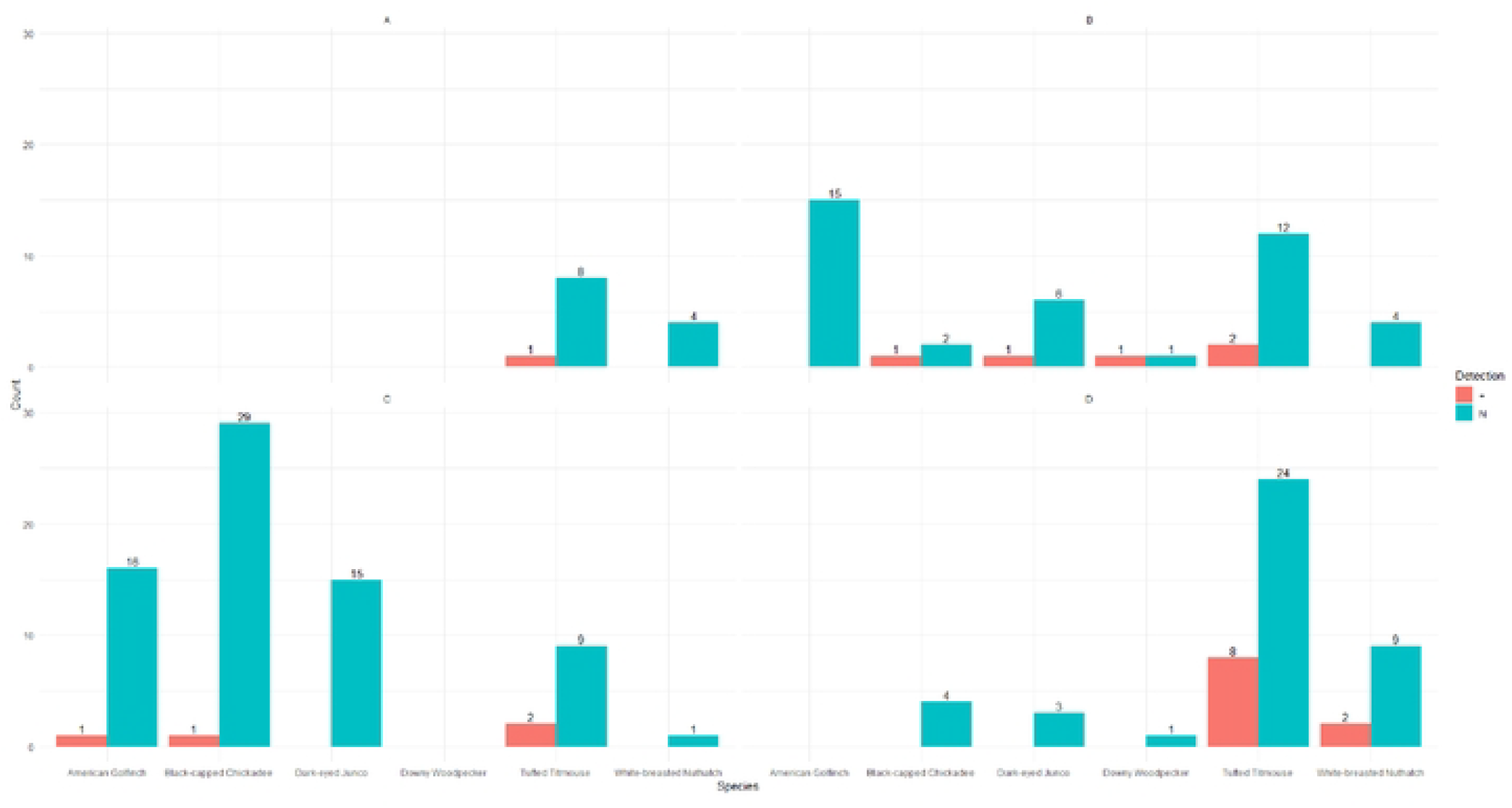
Positive Lcm DNA detections and sample size based on experiment type by species. Sample size (N) is shown in blue and the number of positive detections of Lcm DNA (+) is shown in red. (A) Feather samples, feathers plucked from the breast, head, and vent of birds before any other sampling. (B) Ectoparasite samples, brushing with diatomaceous earth to collect ectoparasites. (C) Bagged bird fecal samples, samples collected out of brown paper bags after birds were kept for > 15 minutes. (D) Caged bird fecal samples, fecal samples from birds kept in caged >4 hours and fed Lcm nematodes.

### Ectoparasite samples

We collected 55 ectoparasite samples from birds. Six samples were positive for Lcm DNA (10.9%). We collected 39 samples at the ANF; one dark-eyed junco had a positive ectoparasite sample. We collected 31 samples at Holden; two black-capped chickadees, two tufted titmice, and one downy woodpecker had positive ectoparasite samples (Fig 4b). We observed no motile nematodes with microscopy.

### Bagged birds’ fecal samples

We collected 90 fecal samples from birds kept in paper bags. Four samples tested positive for Lcm DNA (4.44%). Two came from tufted titmouse, one from an American goldfinch, and one from a black-capped chickadee all at Holden (Fig 4c). We observed no motile nematodes with microscopy.

#### Caged birds’ fecal samples

We collected 59 fecal samples from caged birds. Ten samples were positive for Lcm DNA (16.9%). GenBank BLASTn searches showed that all rDNA sequences were 99% identical to Lcm (MK292138.1). Positive samples came from eight tufted titmice and two white-breasted nuthatches. We caged eleven birds at Holden; two tufted titmice had positive fecal samples. At ANF we caged 31 birds; six tufted titmice and two white-breasted nuthatches had positive fecal samples (Fig 4d). We observed Lcm in fecal matter with microscopy but no motile nematodes.

## Discussion

In this study, we successfully detected the DNA of Lcm nematodes in bird feces, bird ectoparasites, and bird feather samples. These results and methods are important fundamental groundwork for the future of BLD research and understanding and managing its spread. This is among the first studies using DNA to detect plant parasitic nematodes from avian samples. Avian molecular scatology is an emerging field and has been used to detect arthropod, plant, bacterial, and viral DNA [47-51]. We caught all birds in heavily infested BLD areas. If these BLD areas continue to have a high rate of American beech mortality due to beech bark disease and BLD, birds associated with American beech might seek other tree species for food (buds, nuts, insects), nesting, and breeding or overwintering habitat [2,52]. Beech bark disease greatly reduced mature mast-producing American beech trees stimulating the growth of young understory thickets [6]. Young understory American beech thickets have a higher BLD infection rate than mature overstory trees [27]. Birds could acquire Lcm through endozoochory or ectozoochory and spread it to new stands thus spreading BLD.

The bird species we found with Lcm DNA are primarily non-migratory. Tufted-titmouse, black-capped chickadee, and white-breasted nuthatch (Fig 3) were the species with the most positive detections. These three species are common and widespread with population movement occurring primarily from juveniles moving into new territories [53,54]. These species might not be candidates for long-distance dispersal but could still act as local dispersal vectors. These birds can be found in urban and suburban areas, which have a lower detection of BLD compared to more rural areas. As urbanization continues, the edges of these ecotypes become more intertwined and bird communities continue to expand and cooccur in urban areas [55]. American beech in urban areas are often high-value trees in parks and arboreta. Urban trees are more isolated and at risk of encountering tufted titmice, Carolina chickadee (*Poecile carolinensis*), black-capped chickadee, and white-breasted nuthatch.

Most Lcm detections came from tufted titmice in our endozoochory caged experiment. In preliminary trials, we observed red-bellied woodpeckers (*Melanerpes carolinus*), downy woodpeckers, song sparrows (*Melospiza melodia*), black-capped chickadees, tufted titmice, and American goldfinches at bird feeders baited with American beech buds. During our experiment, we detected endozoochory Lcm positives in fecal matter from black-capped chickadees, tufted titmice, and American goldfinches that were caught at feeders filled with infested American beech buds. These positives likely resulted from these birds consuming infested buds and passing detectable nematode DNA. Species observed at feeders baited with infested American beech buds that did not produce positive fecal samples include northern cardinals (*Cardinalis cardinalis*), hairy woodpeckers (*Dryobates villosus*), chipping sparrows (*Spizella passerina*), blue jays (*Cyanocitta cristata*), red-winged blackbirds (*Agelaius phoeniceus*), and house finches. We also caught red-breasted nuthatches (*Sitta canadensis*) and white-throated sparrows (*Zonotrichia albicollis*) but did not collect samples from them. Other insectivorous species, including hooded warbler (*Setophaga citrina*), Acadian flycatcher (*Empidonax virescens*), and brown creeper (*Certhia americana*), were caught and sampled with no positive detections, likely indicating they were bycatch rather than regular visitors to the feeders.

We conducted the bagged bird experiment in 2021 and 2022 before starting the cage bird experiment in 2022 and 2023. Bagged birds had a low positive detection rate (4.4%, n = 90). The lack of direct observation of bagged birds consuming buds could explain the low positive detection. It was difficult to determine if birds were consuming American beech buds, especially buds from trees with BLD. We developed a cage experiment to control and directly observe the birds consuming BLD buds. Using methods primarily intended for long-term confinement and the study of bird’s diet and behavior, we successfully altered them for short-term field application [39]. We knew these caged birds were consuming Lcm, so all should have been positive detections. Detection was higher in caged birds than bagged birds (16.9%, n = 59), but not as high as we predicted, suggesting that Lcm might have difficulty successfully passing through a bird’s digestive tract. When we fed birds buds from BLD trees, rather than feeding them Lcm nematode suspension, we had a higher detection rate. The bud may act as a protective barrier for the nematode during digestion. On one occasion a caged tufted titmouse excreted a partially digested bud, and we found several intact, non-motile, nematodes that tested positive as Lcm.

Soybean and potato cyst nematodes can survive the digestive tract of birds and produce viable eggs [24,25]. Cyst nematodes differ from Lcm greatly, most notably the adult female life stage of cyst nematodes is filled with hundreds of eggs and forms a hard outer cuticle or cyst [56]. This cyst might act as a protective barrier and keep eggs viable while moving through the digestive system. It would have been extremely effort-intensive to isolate only pregnant Lcm females as they are vermiform, much smaller than cysts nematodes, and contain far fewer eggs. We observed all life stages (eggs, juveniles, and mature adults) in nematode suspensions fed to the birds. We successfully detected Lcm DNA in avian fecal samples but never observed Lcm motility. We used microscopy on caged bird fecal samples and observed intact non-motile Lcm nematodes on multiple occasions. The lack of motility does not indicate a lack of viability. We did not use fluorescent markers, which have been used to determine the viability of cyst nematodes [57]. Observing motility and viability of Lcm from avian biological samples, and if those nematodes can cause BLD symptoms on uninfected American beech are important next steps to understanding if birds can act as vectors of BLD. In an ad hoc experiment, we inoculated uninfected American beech seedlings in a greenhouse using methods outlined in Carta et al. [8] with fecal matter from caged birds fed Lcm nematodes. While we did not have any seedlings leaf out with BLD symptoms, we believe continued inoculation experiments are critical for future studies [30]. Understanding the ability of Lcm coming from avian samples to colonize uninfected trees would give valuable insight into the pathogen transmission.

There is limited ongoing research exploring American beech nuts for the presence of Lcm. Over 30 species of mammals and birds consume American beech nuts [2]. White-tailed deer, American black bear, tree squirrels (*Sciurus* spp.), blue jay (*Cyanocitta cristata*), wild turkey (*Meleagris gallopavo*), and ruffed grouse (*Bonasa umbellus*) are all common animals that consume beechnuts. American beech nuts are especially important in the diet of ruffed grouse and affect their habitat use [58,59]. Examining beech nuts for viable Lcm nematodes is an important next step. If viable Lcm exists in beechnuts, ruffed grouse, and other common animals are possible vectors of BLD. We could modify methods used by Servello and Kirkpatrick [60], examining crop contents of hunter-killed ruffed grouse, to look for Lcm nematodes.

Other viable plant pathogen nematode dispersal has been observed with pine wilt disease [61]. Pine sawyer beetles (*Monochamus* spp.) spread pine wilt nematode (*Bursaphelenchus xylophilus*) via motile nematodes moving into the spiracles (external respiratory openings) of beetles, and infesting pine through wounds created during the beetle’s life cycle [62]. Beetles or other insects could be similarly spreading Lcm. Tussock moth caterpillars (*Lymantriinae* spp.) fed American beech leaves with BLD symptoms produced viable Lcm nematodes in their frass (Kantor et al. 2023). Birds could act as a secondary vector through the consumption of caterpillars that have been feeding on BLD leaves. Dark-eyed junco, mountain chickadee (*Poecile gambeli*), and red-breasted nuthatch (*Sitta canadensis*) are known predators of Douglas-fir tussock moth caterpillars (*Orgyia psedotsugata*) and are birds, or closely related to birds, we caught in abundance [63]. Banded tussock moth caterpillars (*Halysidota tessellaris*) have been observed on American beech leaves. Using similar methods to our cage experiment but feeding birds caterpillars that have been fed BLD leaves instead of buds might aid in recovering viable Lcm nematodes from avian feces. Caterpillars might serve as a protective barrier like buds or cysts to keep nematodes viable. Caterpillars are more likely to be consumed by birds than American beech buds. Birds could consume caterpillars feeding on BLD leaves then fly to an uninfected area and excrete viable nematodes. Additionally applying fluorescent stain to non-motile nematodes recovered from fecal matter could help determine viability [64].

While we had a small sample size for feathers, we did not manipulate birds in any way for ectozoochory sampling, unlike the caged bird experiment. It is important to note that ectoparasite and fecal samples from bagged birds often contained feathers. The feather samples were strictly feathers collected before ectoparasite sampling. We collected birds in infested BLD areas thus, nematodes or ectoparasites with nematodes must have been acquired from the environment. In the wake of beech bark disease, heavily foliated American beech thickets have sprouted from the root systems of mature dead American beech [7]. Birds could acquire nematodes directly from leaves on their feathers or from mites with nematodes. Oribatidae mites (*Ommatocepheus clavata*) are associated with American beech and have been observed with live Lcm nematodes on their bodies [65]. Oribatid mites are primarily phytophagous (herbivorous) and other phytophagous mites in the genus *Brevipaplus* have been associated with birds and bird nests [66,67]. *Brevipalpus* mites are vectors of the viral plant pathogen citrus leprosis, initially scaly bark [68]. Though citrus leprosis is a viral pathogen, a similar disease system may occur with birds acquiring Oribatid mites that have Lcm and introducing nematodes to uninfected American beech. Other forest pathogens have been detected environmentally from forest passerines. Forest passerines can acquire viable *Phytophthora ramorum*, the oomycete that causes sudden oak death, in the environment on their feet and feathers [69]. Researchers successfully cultured *P. ramorum* from samples recovered from wild passerines. A limitation of our ectozoochory sampling was the methods were fatal for ectoparasites and we did not observe viability. Using methods by Dadam et al. [69] would be a valuable test to find live, motile, viable Lcm nematodes externally on birds. Testing in August-October, during peak Lcm detection frequency and nematode per leaf sample, would likely provide the highest chance of birds picking up Lcm nematodes environmentally [70].

Ectozoochory appears to be the more likely method of long-distance Lcm dispersal. We did not manipulate birds by placing Lcm nematodes on them externally, thus birds likely acquired Lcm in the environment. Nematodes can travel much farther on birds through ectozoochory than through endozoochory. Through endozoochory, birds have a digestive timeline and can only fly so far before defecation. With ectozoochory, nematodes could attach and remain on a bird indefinitely before being shed and introduced into a new area. Species of feather mites (*Amerodectes* spp.) can survive > 10 days on a host [71]. If we found Lcm nematodes on those mites, migratory birds carrying the mites would be capable of long-distance dispersal. Birds can travel > 200 km/day during migration [72]. If birds are vectors of Lcm, identifying which species are primary vectors could inform predictive models for the future spread of BLD.

## Conclusions

American beech is an important hardwood, hard masting, tree. Beech bark disease has decreased American beech greatly, and American beech continues to be host to numerous other pests. These include beech blight aphid (*Grylloprociphilus imbricator*), beech leaf mining weevil (*Orchestes fagi*), sooty mold (*Scorias spongiosa*), beech erineum mite (*Acalitus fagerinea*), and now beech leaf disease (Lcm). We conducted this experiment to determine the feasibility of birds spreading Lcm. We modified previous methodologies and created new methodologies for detecting nematodes from avian feces, feathers, and ectoparasites. These methods are important for future research of Lcm and BLD in avian vectors but are also broadly applicable to forest pathology, nematology, and ornithology. We successfully detected Lcm DNA from feces, ectoparasites, and feathers. Determining the dispersal vector of Lcm is an important first step in conserving this ecologically and culturally significant tree species.

## Acknowledgments

This research was funded by a grant from the USDA Forest Service, State, Private, and Tribal Forestry (21-DG-11094200-248), in cooperation with West Virginia University accession #1011563, The Holden Arboretum, and The Allegheny National Forest. We thank Mike Watson and Mary Pitts of the Holden Arboretum for their assistance with fieldwork and lab work. We thank Christian Harris of West Virginia University for assistance with fieldwork and lab work. We thank Megan Dudenhoeffer for assistance with molecular methods, editorial comments, and emotional support.

## Notes

### Competing Interest Statement

The authors have declared no competing interest.

